# Spatially resolved diversity in molecular states underlies congenital melanocytic nevi and associated tumors

**DOI:** 10.64898/2026.01.20.700607

**Authors:** Daniel Aldea, Sarah Guégan, Israa Fakih, Mathias Moreno, Arnaud de la Fouchardière, Daniel Pissaloux, Franck Tirode, Joan Anton Puig-Butillé, José-Luis Villanueva-Cañas, Teresa Torres-Moral, Anton Forniés-Mariné, Stéphanie Mallet, Nathalie Degardin, Anne Barlier, Susana Puig, Sylvie Fraitag, Cristina Carrera, Nicolas Macagno, Heather C. Etchevers

## Abstract

The congenital melanocytic nevus (CMN) is a developmental skin disorder characterized by prenatal melanocyte overgrowth, exhibiting heterogeneity in surface size, depth, and clinical behavior. Large/giant CMN carry an elevated, anticipatory melanoma risk relative to more common, small CMN. However, deeper insights into melanocytic states, genomic and epigenomic alterations, and microenvironmental cues governing disease progression are needed to predict lesions disposed to transformation. Although large/giant CMN melanocyte heterogeneity was recently established, spatial organization of these states and their niche interactions are unknown. Using advanced spatial and single-cell transcriptomics and bulk methylomics, we characterized ten CMN, seven CMN-associated proliferative nodules and three clinically diagnosed melanomas arising in CMN from children, integrating data from healthy skin references to contextualize melanocyte states *in situ*. CMN-specific melanocytic states, distinct in differentiation and proliferation, were spatially stratified, with immature melanocytes in deep dermis and more differentiated melanocytes approaching the epidermis. We then constructed a robust atlas of CMN cellular states by integrating single-cell transcriptomic data from five new large/giant CMN with published datasets, using it to deconvolute the spatial information. Cell–cell communication inference uncovered enhanced signaling (e.g. pleiotrophin, IGF1, periostin, semaphorin pathways) between CMN melanocytes, fibroblasts, and hair follicle–associated cells. Analysis of CMN-derived tumors, including longitudinal cases with ≥2 samples, revealed divergent spatial melanocytic transcription distinguishing immune-enriched lesions from tumors with oncogenic/pro-invasive signatures. Collectively, these findings establish a spatially resolved framework linking melanocyte heterogeneity, signaling, and genomic instability in CMN, providing mechanistic insights to refine risk stratification and prognosis for CMN-associated tumors.

## Introduction

The congenital melanocytic nevus (CMN, plural nevi) is a developmental disorder characterized by the extensive prenatal proliferation of melanocytes – the pigment-producing cells of the skin, of neural crest origin – in one or more circumscribed zones of the skin, subcutis, and/or meninges (1). CMN exhibit marked heterogeneity in surface size, depth, and clinical behavior. Isolated, small CMN occur in every 1-2 per hundred births, but are increasingly rare with greater total body surface implicated, with an estimated prevalence of 1/20,000 to 1/500,000 births for the largest forms (2, 3). Children with large (20–40 cm) or giant (>40 cm) (L/G)CMN, based on predicted adult diameter, carry increased lifetime risk of developing proliferative melanocytic neoplasms, sometimes fatal (4, 5). As many as 5% of L/GCMN-bearing children overall may develop cancer, in sun-protected skin or the in central nervous system with similar prevalence (4, 6). Multiple medium CMN are similarly concerning (4).

The genetic and epigenetic regulation of signaling programs within and between cells that govern melanocyte states, spatial organization, and behavior in CMN remain poorly understood. Cell-intrinsic activation of the Mitogen-Activated Protein Kinase (MAPK) pathway, particularly through somatic *NRAS* (p.Q61 to amino acids R, K, or L) or *BRAF* p.V600E variants, accounts for the genetic cause of most, though not all, CMN cases (7). Additional variants, as well as gene fusions or duplications implicating MAPK effectors, have been identified that seem to converge on pathway activation (8, 9).

Although *BRAF* variants are less frequent in the largest CMN than *NRAS*, what little genotype/phenotype correlation exists in large/giant CMN links *BRAF* gain-of-function, often through gene rearrangements, to increased benign nodularity (7, 10, 11), separate or concomitant vascular malformations (12, 13), or proliferative “neurocristic” hamartoma (14). Large and giant CMN often display hamartomic disorganization of skin structures, leading to the current classification that includes clinically ascertainable color heterogeneity, rugosity, nodularity and hairiness (5). However, the mechanisms through which such diverse outcomes can arise in CMN remain unknown.

Spatial context is a critical determinant of melanocyte biology. In normal skin, melanocyte maturation, differentiation, and dendricity are tightly regulated by interactions with keratinocytes, fibroblasts, and other stromal components through paracrine signaling and direct cell–cell contact (15–17). Disruption of these interactions can profoundly alter melanocyte differentiation states and has been implicated in melanoma initiation and progression (18–20). Recent single-cell RNA sequencing (scRNA-seq) studies have revealed transcriptional heterogeneity among melanocytes in three giant CMN (21). However, scRNA-seq approaches inherently lack spatial information in exchange for granularity, limiting insight into how distinct melanocytic states are organized and their developmental trajectories within the complex architecture of the skin. This limitation is particularly relevant in CMN, where melanocytes can be distributed through the entire thickness of the skin, often tracking adnexal structures, vessels, nerves, hypodermal fat and fascia. In these varied locations, they are exposed to ectopic microenvironmental cues that may influence their maturation and functional properties (1).

To overcome this hurdle, in this work we comprehensively mapped melanocyte transcriptional states and intercellular communication networks of large and giant CMN biopsies, as well as an exceptional collection of CMN-associated melanocytic tumors (proliferative nodules and melanoma), using spatial transcriptomics to assess over 15,000 transcriptomes. Furthermore, we analyzed multiple lesions biopsied over a period of years from two children with head CMN, one of whom died from complications of malignant melanoma. We augmented currently available cell atlases at single-cell resolution with five additional, *NRAS*- or *BRAF*-variant large/giant CMN and used these to further annotate and deconvolute the spatial maps. By resolving melanocyte heterogeneity within native tissular contexts and comparing with unaffected control skin, we observed the spatial gradient of melanocyte maturation, identified disease-specific rewiring of signaling pathways, and defined molecular patterns associated with proliferative behavior. Together, our findings provide a spatially informed framework for understanding melanocyte biology in CMN and offer insights into the mechanisms that may govern benign persistence or growth versus malignant transformation in this mosaic RASopathy.

## Results

### Spatial organization and transcriptional identity of melanocytes in congenital melanocytic nevi

Since CMN harbor accumulations of melanocytes not only at the epidermal-dermal junction but also through the thickness of the dermis, and in contrast to so-called blue nevi, are more pigmented toward the surface than at depth (22), we hypothesized that melanocytes followed a maturation gradient from deep to superficial. Spatial transcriptomics (ST) allowed us to test this in an unbiased manner by assessing the transcripts of 18,132 genes, targeted by 53,519 probes, across an array of regularly spaced spots sampling full-thickness skin biopsies from five CMN patients with distinct presentations (Table 1). Representative of the spread of sizes and most typical gene variants, two biopsies were from small, two from medium, and one from a giant CMN from four children and one adult; the two small CMN carried BRAF p.V600E variants, and the others *NRAS* p.Q61K or R (23). To accurately annotate the ST spots, we integrated our dataset with a technically comparable study of unaffected skin samples (24). This control data consisted of five samples of unaffected skin from four donors (33-47 years old). After quality control and integration of all ten sections, 15,466 spots were retained for downstream analysis (Figure 1A).

**Figure 1.**
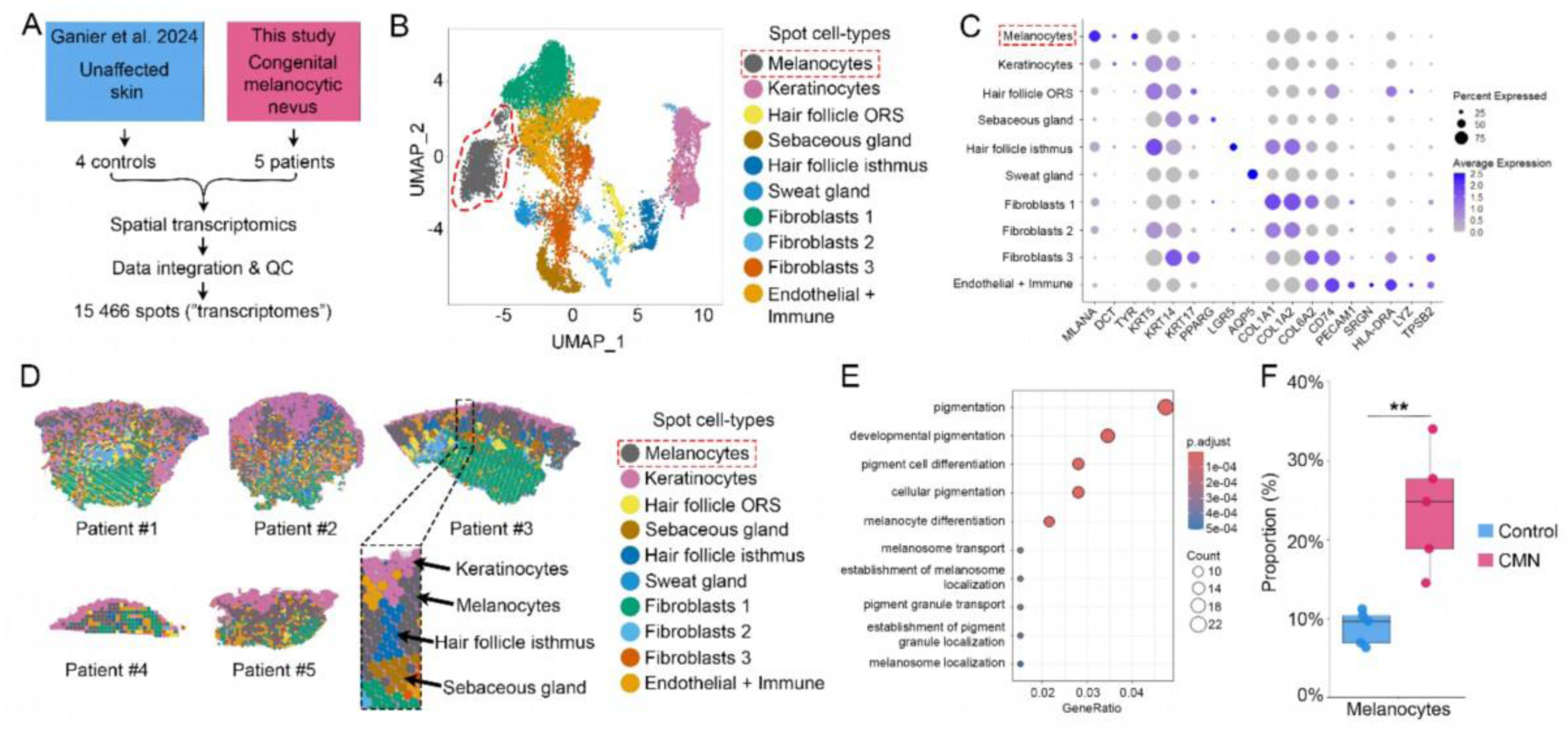
Spatial organization and transcriptional identity of melanocytes in congenital melanocytic nevi. **(A)** Overview of the study design. Skin samples from five patients with congenital melanocytic nevi (CMN) were subjected to spatial transcriptomic profiling and integrated with unaffected skin samples from four control individuals previously published in *Ganier et al.* (24). **(B)** Uniform manifold approximation and projection (UMAP) representation of the integrated dataset following quality control, showing annotation of transcriptomic spots associated with skin cell types. ORS: outer root sheath. **(C)** Dot plot illustrating the expression of canonical skin cell–type marker genes across spot clusters. Dot color intensity indicates the average expression level of each gene within a given cluster, while dot size represents the percentage of spots expressing that gene. Canonical markers were selected based on previously published skin atlases (24, 85). **(D)** Spatial distribution of transcriptomic spot clusters, highlighting the predominant cell types associated with each cluster. **(E)** Gene Ontology (GO) enrichment analysis of differentially expressed genes (DEGs) in the melanocyte-associated cluster. **(F)** Comparative proportions of melanocyte-associated clusters in unaffected skin (Control) and CMN samples. ** P < 0.01, determined using a Wilcoxon rank-sum test.

In an analogous manner to single-cell transcriptomic analysis, we established a UMAP distribution of spot transcriptomes on the basis of most differentially expressed genes and manually annotated the spot clusters with canonical markers of ten dominant skin cell types (Figure 1B; 1C). The spatial distribution of these “spot–cell” types corresponded closely with underlying and histopathological features observed across all samples (Figure 1D). Gene Ontology functional annotation further confirmed that the designated melanocyte cluster was significantly enriched for pigmentation-related biological pathways (Figure 1E; Table 2).

As a necessary validation, CMN tissue displayed a significantly increased number of spots dominated by melanocytes compared with unaffected skin (on average 23.9% vs. 8.8%, respectively), consistent with the true tissue composition (Figure 1F) (1). Together, these results demonstrated strong correspondence between histological features and the spatial organization of transcriptionally defined spot-cell clusters.

### A spatial gradient of melanocyte maturation underlies transcriptional heterogeneity in CMN

Melanocyte transcriptional heterogeneity has been recently shown in CMN patients through a single-cell RNA sequencing (scRNA-seq) study (21). However, the spatial organization of the fine-grained, heterogeneous melanocyte populations has not yet been characterized, nor has their existence been confirmed in an independent cohort. By subclustering the 2,274 melanocyte-dominant spot transcriptomes of the above analysis, we identified three distinct subclusters (Clusters 0, 1, and 2), with Cluster 2 being almost exclusively detected in CMN samples (Figure 2A, 2B). Although Cluster 2 was present in all CMN patients, its abundance varied substantially across individuals #1–5 (Table 1), reflecting the intrinsic cellular heterogeneity of the biopsies (7.8%, 4.6%, 53%, 13.7%, and 16.5%, respectively).

**Figure 2.**
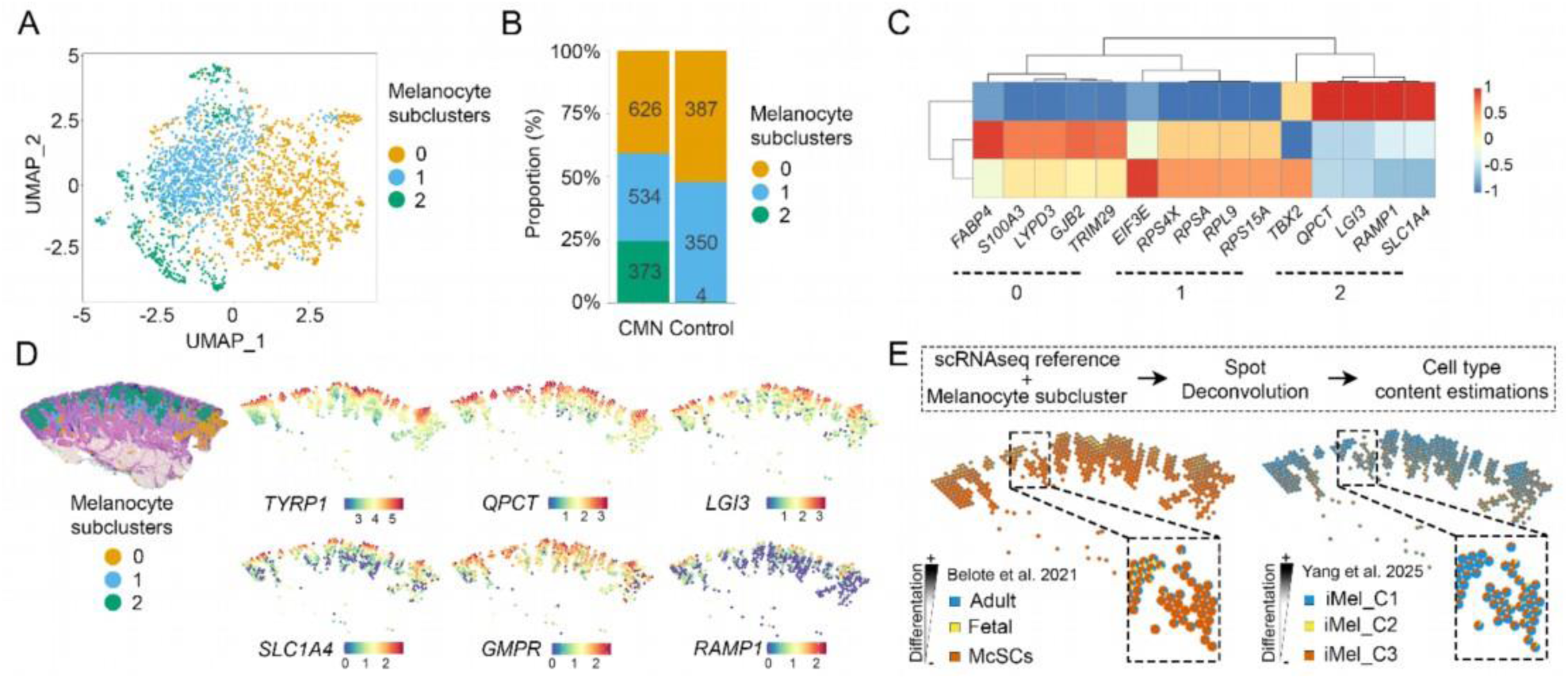
A spatial gradient of melanocyte maturation underlies transcriptional heterogeneity in CMN. **(A)** UMAP representation of melanocyte-associated spots, showing three distinct melanocytic subclusters. **(B)** Relative proportions of each melanocytic subcluster across CMN and control skin samples. **(C)** Heatmap of differentially expressed genes distinguishing the three distinct melanocytic subclusters. **(D)** Representative tissue section from Patient #3 illustrating the spatial distribution of the three melanocytic subclusters, alongside spatial expression patterns of the top differentially expressed genes (DEGs) defining cluster 2. **(E)** Top: Schematic overview of spatial spot deconvolution using reference single-cell atlases from *Belote et al.* and *Yang et al.* (39, 40). Bottom: Representative distribution of estimated melanocytic state composition across spatial transcriptomic spots in Patient #3 for each reference atlas.

Examination of cluster-specific DEGs revealed that cluster 0 was characterized by the expression of *FABP4*, expressed in melanocytes (25); the melanocyte stem cell-specific enzyme *ALDH2* (26) and the functionally related *ALDH3B2*; *GJB2*, a gap-junction protein associated with normal melanocyte development and extracutaneous function (27); and *LYPD3* and *TRIM29*, both of which highly expressed in melanoma (28, 29). The presence of hair follicle-specific keratins (*KRT75, KRT27, KRT71*) testifies to the presence of non-melanocytic cells within cluster 0 spots, borne out by its perifollicular distribution in sections (Figure 2D). Cluster 1 was enriched for multiple ribosomal protein (RP)–related genes, including *RPL9*, *RPS15A*, *RPS4X*, *RPSA,* and *EIF3E*. This signature may reflect elevated protein synthesis, potentially associated with cellular stress responses and cell-cycle arrest (30, 31). *MGP*, encoding the acidic, calcium-binding matrix Gla protein, was modestly if significantly upregulated in this cluster (Figure 2C; Tables 3-4).

In contrast, several genes were transcribed in a conspicuous spatial gradient among the top DEGs defining cluster 2, with stronger expression in more superficial melanocytes (Figure 2C, 2D; Table 5). These include *TYRP1*, a major effector of eumelanin synthesis (32); *QPCT*, encoding glutaminyl-peptide cyclotransferase in melanocytes (33); *LGI3*, a secreted protein associated with pigmentation (34); *SLC1A4*, an amino acid transporter, with tumor-promoting functions (35); *GMPR*, a melanoma invasion suppressor (36) and *RAMP1*, a receptor recently implicated in melanocyte transformation and dedifferentiation (37) (Figure 2D). *TBX2*, recently reported to be expressed in a specific CMN melanocyte subpopulation identified by scRNAseq (21), was differentially expressed in both spatial clusters 1 and 2 relative to cluster 0 (Figure 2C; Table 4).

We hypothesized that this spatial gradient in gene expression reflects distinct states of melanocyte maturation, as melanocytes located deeper in the dermis may not receive keratinocyte-derived maturation signals (15, 16). Consistent with this idea, histopathological studies have described a maturation gradient characterized by stronger expression of HMB-45—a diagnostic marker that recognizes *PMEL*, a melanocyte-specific protein essential for the formation of melanosomes—near the surface, with reduced expression in deeper dermal regions (38).

To identify the contribution of maturation to the spatial gradation of cluster 2 gene expression, we leveraged two published single-cell RNA sequencing reference atlases defining distinct developmental melanocytic states to deconvolute the spatial transcriptomic spots. The atlas from *Belote et al.* comprises melanocytes enriched across multiple body sites, developmental stages, and sexes, whereas the atlas from *Yang et al.* uses melanocytes derived from human pluripotent stem cells to recapitulate melanocyte development *in vitro* (39, 40). This analysis revealed a conspicuous spatial pattern, with less mature melanocytic states from both atlases enriched in deeper dermal regions, while more mature melanocytes were localized closer to the epidermis (Figure 2E).

### Increased intercellular communication in CMN skin

We next applied CellChat to identify signaling pathways underlying the melanocyte maturation gradient and to delineate differences between control and CMN skin. CellChat infers the probability of intercellular communication by integrating gene expression profiles and spatial proximity with prior knowledge of ligand–receptor interactions (41). We observed a significant increase in inferred cell–cell communication and interaction strength in CMN compared with control skin (number of interactions: 3,580 vs. 928, respectively) (Figure 3A-B). Melanocytes, Fibroblasts 1, Fibroblasts 2, and HF_isthmus clusters displayed the most sizeable changes in both signal sending and receiving in CMN compared to control samples (Figure 3C).

**Figure 3.**
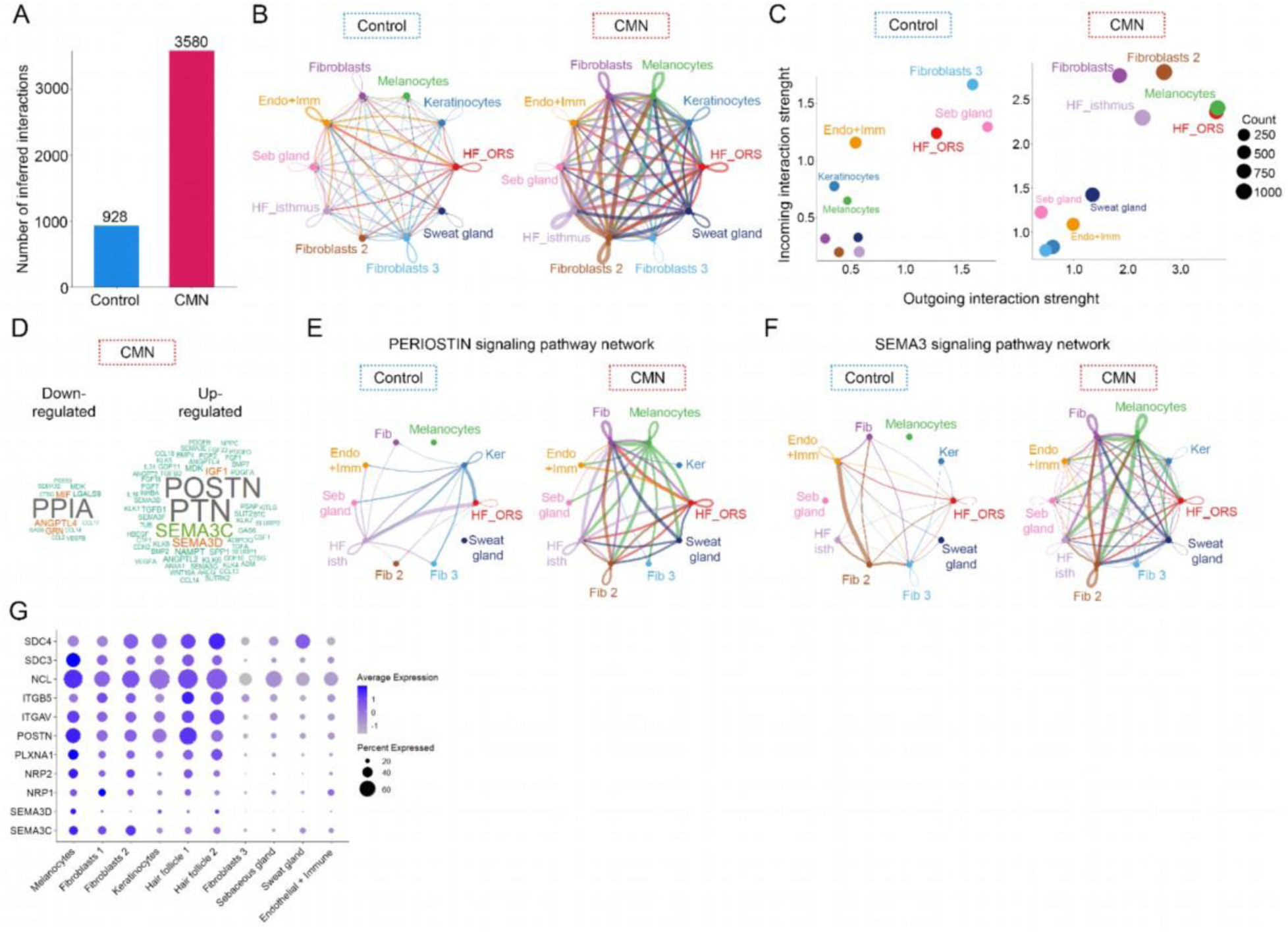
Increased intercellular communication in CMN skin. **(A)** Total number of inferred ligand–receptor interactions in control versus CMN samples. **(B)** Circle plots depicting inferred ligand–receptor interactions among skin cell–type clusters in control versus CMN samples; edge width represents the number of predicted interactions between clusters. **(C)** Scatter plots showing the major cell-type sources contributing to outgoing (sending) and incoming (receiving) signaling in control and CMN samples. **(D)** Word clouds illustrating ligands significantly upregulated or downregulated in CMN compared with control skin. **(E-F)** Circle plots showing the interactions among different cell types of PERIOSTIN and SEMA3 signaling pathways. **(G)** Dot plot showing expression levels of genes involved in the identified ligand–receptor interactions across skin cell–type clusters. Dot color intensity indicates average gene expression within each cluster, while dot size represents the proportion of spots expressing the gene.

We next investigated signaling pathways flagged by CellChat as differentially regulated in CMN compared with control skin. Consistent with previous findings, pleiotrophin (PTN) and insulin-like growth factor-1 (IGF1) signaling pathways were significantly upregulated in CMN samples relative to controls (Figure 3C)(21). PTN is a cytokine with diverse functions, including the growth of glial progenitor cells, promotion of neurite outgrowth, and regulation of angiogenesis (42), whereas IGF1 is a potent growth factor with established roles in cell survival, differentiation, and tissue development (43). Early bulk transcriptomes of undifferentiated, primary human neural crest cells in culture also show significant IGF1 pathway upregulation (44). Furthermore, we identified increased activation of periostin (POSTN) and semaphorin (SEMA3C and SEMA3D) signaling pathways (Figure 3D-F). Both POSTN and semaphorins are involved in tissue remodeling, cell migration, and cell–cell communication. Furthermore, melanoma cells have been shown as sources of POSTN, and several members of the semaphorin family have been implicated in the regulation of melanocyte dendrite formation (45–49).

Analysis of inferred ligand–receptor pairs revealed that the most significant interactions included POSTN–(ITGAV+ITGB5), PTN–NCL, PTN–SDC3, PTN–SDC4, SEMA3C/D–PLXND1, and SEMA3C/D–(NRP1/2–PLXNA1) (Figure 3G). These ligands and receptors were highly expressed in Melanocytes, Fibroblast 1, Fibroblast 2, and HF isthmus clusters (Figure 3F).

Cell–cell communication between keratinocytes and melanocytes is well recognized as essential for melanocyte maturation (15, 16). Although keratinocyte–melanocyte interactions were not among the major contributors to overall outgoing (sending) or incoming (receiving) signaling in either control or CMN skin (Figure 3C), we nevertheless observed increased inferred signaling through the PTN, SEMA3C/D, and IGF1 pathways within keratinocyte–melanocyte interactions in CMN, consistent with the overall upregulation of these signaling pathways in CMN tissues.

### Divergent spatial and transcriptional melanocytic programs in L/GCMN-associated atypical tumors

Patients with large or giant CMN (L/GCMN) may develop proliferative nodules, intermediate neoplasms such as melanocytomas, or superficial atypical proliferations. In most cases, these lesions are benign or self-limiting; however, in rare instances, some can progress to melanoma (50, 51). Proliferative nodules, atypical proliferative nodules, and dermal melanomas arising within L/GCMN may require ancillary techniques for a formal diagnosis, including the assessment of 5-hmC, loss of H3K27me3 expression, and, frequently, genomic profiling (52–54). However, the molecular mechanisms driving the transition from nevus melanocytes to intermediate neoplasms and, ultimately, melanoma remain poorly defined, particularly with respect to their spatial organization and functional interactions with the surrounding nevus tissue.

To uncover molecular and signaling pathways dysregulated in this context, we performed ST profiling of additional samples from children bearing CMN-associated tumors (Patients #6 and #7). Histopathological evaluation classified Tumors #1 and #3 as atypical and potentially malignant, whereas Tumor #2 displayed features consistent with a benign proliferative nodular (PN) lesion. These atypical tumors harbored distinct genetic alterations, including a PKC fusion (t(*PTPRJ*;*PRKCA*)) and *BRAF* mutations. The clinical and molecular characteristics of these patients are summarized in Table 1.

Following quality control and integration of all fourteen samples, including CMN and control skin samples, a total of 24,504 spatial transcriptomic spots were retained. Of these, 10,591 spots were annotated as melanocyte-dominant and were used for downstream analyses (Figure 4A).

**Figure 4.**
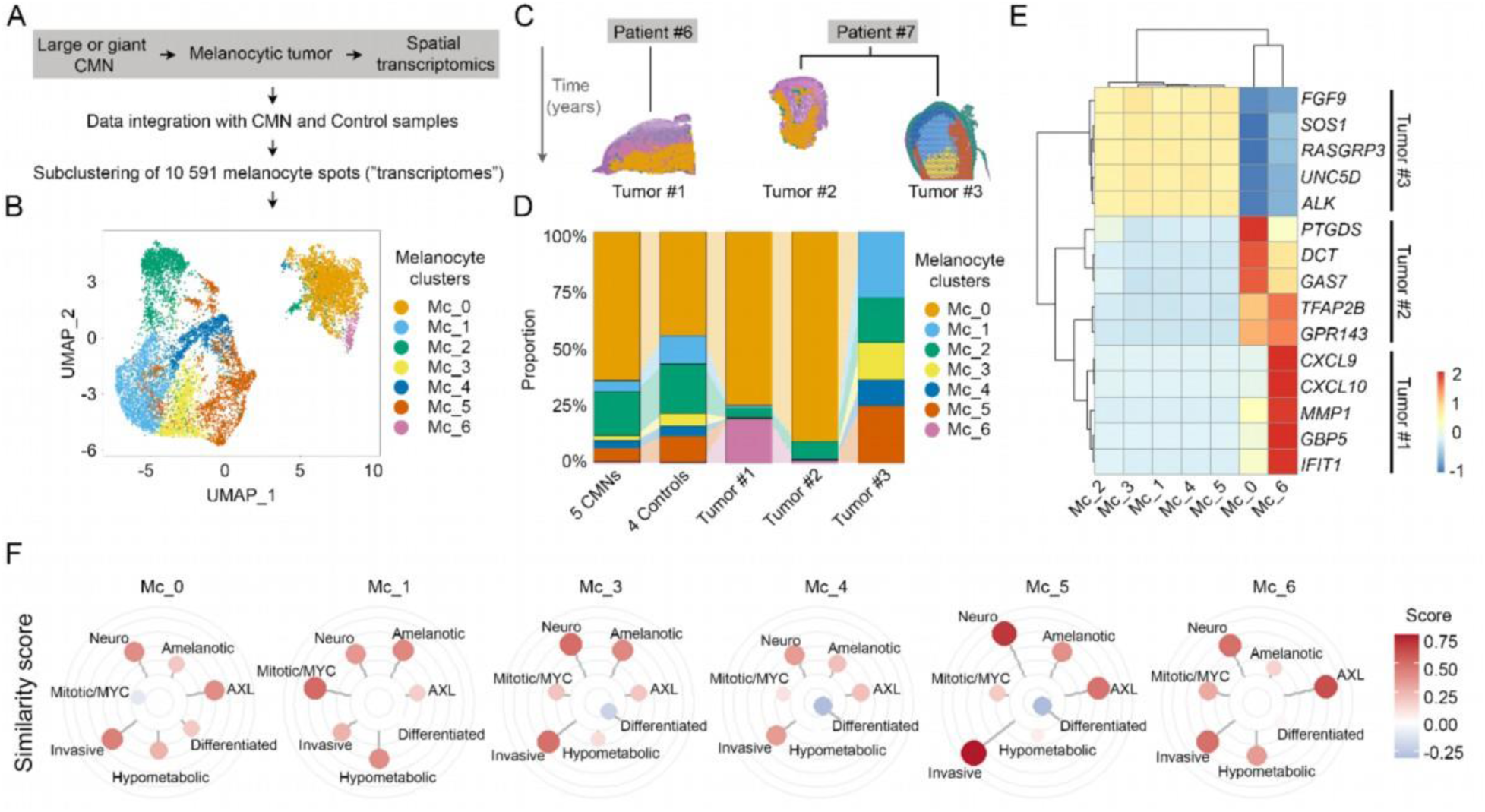
Divergent spatial and transcriptional melanocytic programs in L/GCMN-associated atypical tumors. **(A)** Samples from two patients with atypical melanocytic tumors were subjected to spatial transcriptomic profiling and integrated with CMN and control skin samples. **(B)** UMAP representation showing seven distinct melanocytic clusters identified after subclustering the integrated dataset. **(C)** Spatial distribution of melanocytic clusters across the two patients and the three tumors sampled over time. **(D)** Relative proportions of the seven melanocytic subclusters across tumors, CMN, and control skin samples. **(E)** Heatmap of differentially expressed genes distinguishing Tumor #1 (enriched for Mc_6), Tumor #2 (enriched for Mc_0), and Tumor #3 (enriched for Mc_1, Mc_3, Mc_4, and Mc_5). **(F)** Radial plots showing enrichment scores of melanocytic transcriptional signatures defined by *Hu et al.* (70) across the distinct melanocytic subclusters (Mc_0, Mc_1, Mc_3, Mc_4, Mc_5 and Mc_6).

Among these, we identified seven distinct melanocytic clusters (Mc_0–Mc_6) (Figure 4B), whose spatial distributions varied markedly across the tumors analyzed (Figure 4C). Mc_2 was consistently detected across all samples. Tumor #1 was predominantly composed of Mc_0, a cluster shared with both CMN and control skin, and also showed marked enrichment in Mc_6, which accounted for 75% and 19.3% of melanocytic spots within Tumor #1, respectively (Figure 4D).

Intriguingly, although Tumor #2 and Tumor #3 were derived from the same patient, they displayed distinct transcriptional and spatial cluster distributions. Tumor #2 consisted almost exclusively of Mc_0, which presented 90.5% of melanocytic spots. In contrast, Tumor #3 showed an increased proportion of Mc_1, Mc_3, Mc_4, and Mc_6, with enrichments of 3.3-, 4.4-, 2.9-, and 2.9-fold, respectively, compared with CMN and normal skin samples combined (Figure 4D).

We next examined transcriptional signatures of the melanocytic clusters enriched in each tumor. Tumor #1 (enriched in Mc_6) showed an increase in genes associated with immune activation, regulation of the tumor microenvironment, and cancer progression, including *CXCL9*, *CXCL10*, *GBP5*, *MMP1*, and *IFIT1* (55–59), suggesting an inflammatory and microenvironmentally active tumor state. Tumor #2 (enriched in Mc_0) displayed a transcriptional signature largely shared with CMN and control skin, characterized by elevated expression of lineage-specific melanocytic genes such as *DCT*, *TFAP2B*, and *GPR143* (60–62), as well as *PTGDS*, which is implicated in the regulation of melanocyte proliferation. This profile is consistent with a standard CMN melanocytic state, in contact with *FLG1/2*- and *KRT1/5*-expressing keratinocytes. In contrast, Tumor #3 (enriched in Mc_1, Mc_3, Mc_4, and Mc_5) was characterized by increased expression of genes frequently dysregulated in cancer, including *UNC5D*, *ALK*, *SOS1*, *FGF9*, and *RASGRP3* (65–69), indicative of a transcriptional program associated with oncogenic signaling and potentially increased malignant potential (Figure 4E; Tables 6-8).

To further define the melanocytic programs characterizing each subcluster, we applied the classifier developed by Hu et al., which integrates 39 published melanocyte gene signatures (70). Mc_0 was the only cluster exhibiting a differentiated melanocytic program, whereas Mc_1 was dominated by mitotic and MYC-associated signatures. Mc_3 and Mc_4 showed enrichment for neural crest–like and invasive profiles. Mc_5 displayed the strongest invasive and neural crest–like signatures, while Mc_6 exhibited the highest enrichment for *AXL*-driven states, together with marked neural crest–like and invasive signatures, consistent with a less differentiated melanocytic state (Figure 4F and Tables 9-14).

Together, these data highlight two distinct tumor signaling states. Tumors #1 and #3 were characterized by transcriptional and signaling programs associated with immune engagement, microenvironmental remodeling, and features linked to increased tumor aggressiveness. In contrast, Tumor #2, which was less atypical and classified as benign, retained a melanocytic program closely resembling that observed in CMN and normal skin.

## Discussion

In this study, we established a spatially resolved framework of melanocyte heterogeneity in a broad range of CMN presentations, showing how distinct melanocytic states are organized within the skin microenvironment. A transcriptomic maturation gradient in CMN melanocytes exists that was previously unsuspected in normal skin, with less differentiated cells residing in deeper dermal compartments and more mature melanocytes positioned near the epidermis in proximity to keratinocytes. These findings highlight spatial context as a critical correlate to melanocytic phenotype.

In normal skin, melanocyte differentiation and homeostasis are tightly regulated by keratinocyte-derived signals, including growth factors and cell–cell contact–mediated cues that promote dendricity, terminal differentiation, and pigment production (15–17). Our data suggest that melanocytes residing deeper in the dermis, where contact with keratinocytes is limited or absent, may escape these maturation signals and instead retain transcriptional programs associated with less differentiated states. Such spatially constrained differentiation is likely particularly relevant in CMN, where both pigmented and unpigmented melanocytes are distributed throughout the dermis and adnexal structures. This organization establishes a network of signals—including autocrine, paracrine, and extracellular matrix–mediated cues—that collectively influence melanocyte behavior. Permissive niches may support melanocyte survival and expansion while maintaining a pool of less differentiated cells with increased plasticity (39, 71). One indication of relevance in the context of L/GCMN is after repeated plastic surgeries or the less common techniques of curettage/dermabrasion, residual unpigmented melanocytes in the deeper dermis assume a more superficial relative position to the healing epidermis and can differentiate into cells that renew the pigmentation of the original lesional area or “bleed-through” into adjacent or burgeoning underlying fibrotic scar (72, 73). Likewise, in hypopigmented burn scar tissue, resident unpigmented (and thus invisible) melanocytes can be stimulated to repigment with alpha-melanocyte stimulating hormone that is normally supplied by keratinocytes; the major difference between hypo- and hyperpigmented scars is melanocyte dendricity rather than number (74).

Among novel signaling activated in intact CMN, were semaphorin, pleiotrophin, IGF-1, and periostin-mediated pathways. Semaphorins, including SEMA3C and SEMA3D, are well-established regulators of neural crest cell (NCC) migration during development (47, 75, 76). In melanocytes, which are derived from NCCs, semaphorins have been shown to regulate dendrite formation and interactions with keratinocytes, while in melanoma, they exhibit context-dependent roles in invasion and metastasis (45, 48, 77). SEMA3C has been shown to drive epithelial-to-mesenchymal transition and promote stem-like characteristics in multiple tumor types (78–80). Thus, increased SEMA3C transcription in CMN may contribute to reduced terminal differentiation and sustained plasticity in melanocytes.

Our analysis of CMN-associated atypical tumors reveals marked spatial and transcriptional heterogeneity. Tumors arising in the context of L/GCMN exhibited distinct melanocytic programs, including CMN-like, invasive, neural crest–like, and AXL-driven phenotypes. Cellular heterogeneity is a hallmark of melanoma and correlates with worse prognosis, as increased intratumoral diversity promotes rapid tumor growth, metastasis, and resistance to treatment (81–84). In this context, identifying dysregulated gene panels and transcriptional programs in CMN-associated tumors may yield valuable biomarkers for improved risk stratification and prognostic assessment in affected patients.

This integrated, spatially informed framework provides new insight into how most CMN remain benign despite widespread oncogenic mutations, while a subset acquires the competence to progress toward malignancy. More broadly, our findings underscore the importance of tissue architecture and cell–cell communication in shaping melanocyte fate and tumor risk, with potential implications for risk stratification and therapeutic targeting in CMN-associated neoplasia.

### Limitations of the study

This study has several limitations. While spatial transcriptomics preserves tissue architecture, the resolution of the Visium platform captures multiple cells per spot, limiting the ability to assign transcriptional programs to individual cells and to fully disentangle cell–cell interactions at single-cell resolution. The relatively limited number of CMN-associated tumor samples, although the fruit of years of multicentric recruitment, reflects the rarity of these lesions and may constrain the generalizability of our findings. Future studies integrating higher-resolution spatial approaches with bulk and single-cell genomic profiling will be necessary to further refine the relationship between spatial organization, genomic instability, and malignant progression in CMN.

## Acknowledgments

We are grateful to C. Humbert, C. Castro, and C. Aubert of the MMG Genomics and Bioinformatics facility (GBiM) for assistance with spatial transcriptomics and analyses.

## Author Contributions

D.A. wrote the article with the input of H.C.E. and N.M. D.A., M.M., and I.F., performed the experiments. S.G. and J.A.P.B. supplied patient metadata and transcriptomics data. D.A., D. P., F. T., A.F.M., J.L.V.C. and H.C.E. performed data analyses. S.F., N.D., A. de la F., S.M., T. T.M., N.M. and C.C., provided patient metadata and samples. S.G., A.B., S. P., N. M. and H.C.E. obtained funding for this work. D.A., N.M., and H.C.E. analyzed and interpreted the data. N.M. and H.C.E. supervised the experiments and revised the article.

## Materials and Methods

### Patients

Samples from five children affected with congenital melanocytic nevi were taken after independently scheduled resection in reconstructive surgery with written, informed parental consent. This study was carried out under the aegis of the nationally appointed ethical review board, CPP, under authorization 214 C03. The histopathological samples were provided by the Assistance Publique Hôpitaux de Marseille, Biological Resources Center (BRC AP-HM Biobank), CRB-TBM component (NF S96-900 & ISO 9001 v2015 Certification). The resources used belong to a biological sample collection declared to the French Ministry of Health (Declaration: DC-2013-1781), whose use for research purposes was authorized by the French Ministry of Higher Education, Research and Innovation (Authorization: AC-2011-2018-3105).

### Visium spatial transcriptomic assays

#### Preparation of Visium spatial gene expression libraries

Formalin-fixed, paraffin-embedded (FFPE) skin tissues were sectioned at a thickness of 10 μm and mounted onto slides from the Visium Spatial Gene Expression Slide & Reagent Kit (10x Genomics), which contains 55 μm capture spots arranged at 100 μm center-to-center spacing. Spatial gene expression libraries were prepared according to the manufacturer’s instructions. Before sequencing, tissue sections were stained with hematoxylin and eosin (H&E), and images were acquired at 20X magnification using an Axioscan Z1 scanner (Zeiss, MMG Imaging Platform).

#### Sequencing and data processing

Final libraries were sequenced on a NovaSeq 6000 system at the GBiM facility using the following configuration: Read 1, 28 cycles; i5 index, 10 cycles; i7 index, 10 cycles; Read 2, 50 cycles. Between 30 and 90 million raw reads were generated per sample. After quality control, reads were aligned and processed using Space Ranger v2.0.1 (10x Genomics) to generate a unique molecular identifier (UMI) count matrix for each spatial spot. Tissue-covered spots were identified from the H&E images using the manual alignment tool in Loupe Browser v6.4.1 (10x Genomics), and only tissue-associated barcodes were retained for downstream analyses.

#### Analysis, visualization, and integration of spatial transcriptomic data

After quality control filtering (nCount_Spatial > 200 and nCount_Spatial < 50,000), spatial transcriptomic data were analyzed using Seurat v5.2.0 (86). The R package SCTransform was used for normalization and variance stabilization (87). After inspection and comparison of individual tissue sections, datasets were integrated using IntegrateLayers (method = HarmonyIntegration) implemented in Seurat v5.2.0. Integrated spots were clustered using 30 dimensions at a resolution of 0.3, and clusters were manually annotated based on canonical skin cell markers. CMN patient samples (n = 5) were integrated with a control group consisting of unaffected skin from four participants younger than 50 years (five samples in total)(24).

#### Differential expression analyses, gene ontology enrichment, and melanocytic signature

Differentially expressed genes (DEGs) among clusters were identified using the Seurat function *FindAllMarkers*. For Figures 1 and 2, genes with a log2 fold change greater than 1, an adjusted p-value controlling for false discovery rate (FDR < 0.05), and expression in at least 25% of spots in the first condition were considered differentially expressed. For Figure 4, the fold-change thresholds used are specified in the corresponding tables. Identified DEGs were used as input for Gene Ontology analysis or for melanocytic signature assessment.

Gene Ontology enrichment analysis was performed using the *enrichGO* function from the clusterProfiler R package with default parameters, focusing on Biological Process terms. Enrichment significance was assessed using a hypergeometric test, and p-values were adjusted for multiple testing using the Benjamini–Hochberg method. Enriched GO terms were visualized using the *dotplot* function from clusterProfiler (88, 89).

Melanocytic transcriptional signatures were assessed using the WIMMS (What Is My Melanocytic Signature) Shiny application developed by Hu et al. (https://wimms.tanlab.org/) (70). DEGs from the tested clusters were used as input (Tables 7-12). Radial plots were generated using the ggplot2 R package.

#### Spatial transcriptomic data deconvolution

Redeconve was used to estimate the cellular composition of spatial transcriptomic spots (90). Two publicly available reference atlases were used for deconvolution: the dataset from *Belote et al.*, which comprises melanocytes enriched across multiple body sites, developmental stages, and sexes, and the atlas from *Yang et al.*, which uses melanocytes derived from human pluripotent stem cells to recapitulate melanocyte development *in vitro* (39, 40). Redeconve was run using default parameters, and results were visualized as pie charts for each spatial spot and projected onto the spatial transcriptomic tissue section images to illustrate the estimated proportions of each cell type per spot.

#### Cell–cell communication analysis

Inference and visualization of cell–cell crosstalk among distinct skin cell types were performed using the CellChat v2 R package, which enables the analysis of spatially resolved transcriptomics datasets (41). CellChat incorporates a comprehensive database of ligand–receptor interactions, including soluble agonists and antagonists as well as stimulatory and inhibitory membrane-bound co-receptors, to infer cell–cell communication networks and spatial interaction distances. CellChat was run using default parameters, and after inspection of inferred cell–cell communication interactions in individual samples, comparative analyses were performed by integrating data from all four unaffected skin samples and five CMN patient samples described above.

### Declaration of generative AI and AI-assisted technologies in the writing process

During the preparation of this manuscript, the authors used AI-assisted tools (ChatGPT, Mistral) solely to improve language and clarity through copy-editing. No content was generated by AI. All text was reviewed and edited by the authors, who take full responsibility for the final manuscript.

## Data availability

All data are available in the manuscript or the supplementary materials. Methylome, single-cell transcriptome, and spatial transcriptome data are deposited on GEO, subject to embargo until publication.

## References

1. Maher NG, Scolyer RA, Colebatch AJ. Biology and genetics of acquired and congenital melanocytic naevi. Pathology. 2023;55(2):169–177.

2. Alper JC, Holmes LB. The incidence and significance of birthmarks in a cohort of 4,641 newborns. Pediatr Dermatol. 1983;1(1):58–68.

3. Castilla EE, da Graça Dutra M, Orioli-Parreiras IM. Epidemiology of congenital pigmented naevi: I. Incidence rates and relative frequencies. Br J Dermatol. 1981;104(3):307–315.

4. Kinsler VA, et al. Melanoma in congenital melanocytic naevi. Br J Dermatol. 2017;176(5):1131–1143.

5. Krengel S, et al. New recommendations for the categorization of cutaneous features of congenital melanocytic nevi. J Am Acad Dermatol. 2013;68(3):441–451.

6. Krengel S, Hauschild A, Schäfer T. Melanoma risk in congenital melanocytic naevi: a systematic review. Br J Dermatol. 2006;155(1):1–8.

7. Salgado CM, et al. BRAF Mutations are Also Associated with Neurocutaneous Melanocytosis and Large/Giant Congenital Melanocytic Nevi. Pediatr Dev Pathol. 2015;18(1):1–9.

8. Martins da Silva V, et al. Genetic Abnormalities in Large to Giant Congenital Nevi: Beyond NRAS Mutations. J Invest Dermatol. 2019;139(4):900–908.

9. de la Fouchardière A, et al. Histologic and Genetic Features of 51 Melanocytic Neoplasms With Protein Kinase C Fusion Genes. Mod Pathol. 2023;36(11):100286.

10. Martin SB, et al. Mosaic BRAF Fusions Are a Recurrent Cause of Congenital Melanocytic Nevi Targetable by MAPK Pathway Inhibition. Journal of Investigative Dermatology. 2024;144(3):593–600.e7.

11. Dessars B, et al. Chromosomal translocations as a mechanism of BRAF activation in two cases of large congenital melanocytic nevi. The Journal of Investigative Dermatology. 2007;127(6):1468–70.

12. A l-Olabi L, et al. Mosaic RAS/MAPK variants cause sporadic vascular malformations which respond to targeted therapy. The Journal of Clinical Investigation. 2018;128(4):1496–1508.

13. Etchevers HC, et al. Giant congenital melanocytic nevus with vascular malformation and epidermal cysts associated with a somatic activating mutation in BRAF. Pigment Cell Melanoma Res. 2018;31(3):437–441.

14. El -Rayes D, et al. Congenital Melanocytic Nevus with Neurocristic Cutaneous Hamartoma: A Case Report. Dermatopathology. 2025;12(2):12.

15. Prospéri M-T, et al. Extracellular vesicles released by keratinocytes regulate melanosome maturation, melanocyte dendricity, and pigment transfer. Proc Natl Acad Sci U S A. 2024;121(16):e2321323121.

16. Hirobe T. Keratinocytes regulate the function of melanocytes. Dermatologica Sinica. 2014;32(4):200–204.

17. Klar AS, et al. Human Adipose Mesenchymal Cells Inhibit Melanocyte Differentiation and the Pigmentation of Human Skin via Increased Expression of TGF-β1. Journal of Investigative Dermatology. 2017;137(12):2560–2569.

18. Mazurkiewicz J, et al. Stromal Cells Present in the Melanoma Niche Affect Tumor Invasiveness and Its Resistance to Therapy. Int J Mol Sci. 2021;22(2):529.

19. Ju RJ, Stehbens SJ, Haass NK. The Role of Melanoma Cell-Stroma Interaction in Cell Motility, Invasion, and Metastasis. Front Med. 2018;5. 10.3389/fmed.2018.00307.

20. Marrapodi R, et al. Melanoma–Keratinocyte Crosstalk Participates in Melanoma Progression with Mechanisms Partially Overlapping with Those of Cancer-Associated Fibroblasts. International Journal of Molecular Sciences. 2025;26(16). 10.3390/ijms26167901.

21. Tan W, et al. Single-cell RNA sequencing reveals transcriptomic landscape and potential targets for large/giant congenital melanocytic nevi. Br J Dermatol. 2025;ljaf360.

22. Li Y, et al. Diagnostic Utility of Molecular Profiling in a Giant Cellular Blue Nevus. Am J Dermatopathol. [published online ahead of print: November 24, 2025]. 10.1097/DAD.0000000000003176.

23. Ichi i-Nakato N, et al. High frequency of BRAFV600E mutation in acquired nevi and small congenital nevi, but low frequency of mutation in medium-sized congenital nevi. J Invest Dermatol. 2006;126(9):2111–2118.

24. Ganier C, et al. Multiscale spatial mapping of cell populations across anatomical sites in healthy human skin and basal cell carcinoma. Proc Natl Acad Sci U S A. 2024;121(2):e2313326120.

25. Elmasri H, et al. Fatty acid binding protein 4 is a target of VEGF and a regulator of cell proliferation in endothelial cells. FASEB J. 2009;23(11):3865–3873.

26. Brunsdon H, et al. Aldh2 is a lineage-specific metabolic gatekeeper in melanocyte stem cells. Development. 2022;149(10):dev200277.

27. Guo J, et al. GJB2 gene therapy and conditional deletion reveal developmental stage-dependent effects on inner ear structure and function. Mol Ther Methods Clin Dev. 2021;23:319–333.

28. Hu Y, et al. LY6/PLAUR domain containing 3 (LYPD3) maintains melanoma cell stemness and mediates an immunosuppressive microenvironment. Biol Direct. 2023;18:72.

29. Xia Y, Zhao J, Yang C. Identification of key genes and pathways for melanoma in the TRIM family. Cancer Med. 2020;9(23):8989–9005.

30. Kang J, et al. Ribosomal proteins and human diseases: molecular mechanisms and targeted therapy. Sig Transduct Target Ther. 2021;6(1):323.

31. Sharma DK, et al. Role of Eukaryotic Initiation Factors during Cellular Stress and Cancer Progression. J Nucleic Acids. 2016;2016:8235121.

32. Gautron A, et al. Human TYRP1: Two functions for a single gene? Pigment Cell & Melanoma Research. 2021;34(5):836–852.

33. Yin L, et al. Identification of Genes Expressed in Hyperpigmented Skin using Meta-Analysis of Microarray Datasets. J Invest Dermatol. 2015;135(10):2455–2463.

34. Farag AGA, et al. Leucine-rich glioma inactivated 3: a novel keratinocyte-derived melanogenic cytokine in vitiligo patients. An Bras Dermatol. 2019;94(4):434–441.

35. Hushmandi K, et al. Amino acid transporters within the solute carrier superfamily: Underappreciated proteins and novel opportunities for cancer therapy. Molecular Metabolism. 2024;84:101952.

36. Wawrzyniak JA, et al. A Purine Nucleotide Biosynthesis Enzyme Guanosine Monophosphate Reductase is a Suppressor of Melanoma Invasion. Cell Rep. 2013;5(2):493–507.

37. McGrail K, et al. Transcriptional reprogramming triggered by neonatal UV radiation or Lkb1 loss prevents BRAFV600E-induced growth arrest in melanocytes. Oncogene. 2025;44(21):1592–1608.

38. Rasic D, et al. Diagnostic utility of combining PRAME and HMB-45 stains in primary melanocytic tumors. Annals of Diagnostic Pathology. 2023;67:152211.

39. Yang J, et al. Insights into human melanocyte development and characteristics through pluripotent stem cells combined with single-cell sequencing. iScience. 2025;28(5):112373.

40. Belote RL, et al. Human melanocyte development and melanoma dedifferentiation at single-cell resolution. Nat Cell Biol. 2021;23(9):1035–1047.

41. Jin S, Plikus MV, Nie Q. CellChat for systematic analysis of cell–cell communication from single-cell transcriptomics. Nat Protoc. 2025;20(1):180–219.

42. Deuel TF, et al. Pleiotrophin: a cytokine with diverse functions and a novel signaling pathway. Arch Biochem Biophys. 2002;397(2):162–171.

43. Werner H. The IGF1 Signaling Pathway: From Basic Concepts to Therapeutic Opportunities. Int J Mol Sci. 2023;24(19):14882.

44. Thomas S, et al. Human neural crest cells display molecular and phenotypic hallmarks of stem cells. Hum Mol Genet. 2008;17(21):3411–3425.

45. Scott GA, McClelland LA, Fricke AF. Semaphorin 7a promotes spreading and dendricity in human melanocytes through beta1-integrins. J Invest Dermatol. 2008;128(1):151–161.

46. Wang Z, et al. Periostin: an emerging activator of multiple signaling pathways. J Cell Commun Signal. 2022;16(4):515–530.

47. Alto LT, Terman JR. Semaphorins and their Signaling Mechanisms. Methods Mol Biol. 2017;1493:1–25.

48. Li T, et al. Regenerative Hair Pigmentation via Skin Organoids: Adaptive Patterning Mediated by Collagen VI and Semaphorin 3C. Adv Sci (Weinh). 2025;12(36):e02436.

49. Tilman G, et al. Human periostin gene expression in normal tissues, tumors and melanoma: evidences for periostin production by both stromal and melanoma cells. Mol Cancer. 2007;6:80.

50. Salgado CM, Tomás-Velázquez A, Reyes-Múgica M. Congenital melanocytic neoplasms: clinical, histopathological and recent molecular developments. Virchows Arch. 2025;486(1):165–176.

51. Scard C, et al. Risk of melanoma in congenital melanocytic nevi of all sizes: A systematic review. Journal of the European Academy of Dermatology and Venereology. 2023;37(1):32–39.

52. Macagno N, et al. Reduced H3K27me3 Expression is Common in Nodular Melanomas of Childhood Associated With Congenital Melanocytic Nevi But Not in Proliferative Nodules. Am J Surg Pathol. 2018;42(5):701–704.

53. Busam KJ, et al. Reduced H3K27me3 Expression is Common in Nodular Melanomas of Childhood Associated with Congenital Melanocytic Nevi but Not in Proliferative Nodules. American Journal of Surgical Pathology. 2017;41(3):396–404.

54. Miele E, et al. Epigenomic characterization and therapeutic challenges of melanoma arising in giant nevi in pediatric patients. Discov Oncol. 2025;16:2164.

55. Tokunaga R, et al. CXCL9, CXCL10, CXCL11/CXCR3 axis for immune activation - a target for novel cancer therapy. Cancer Treat Rev. 2018;63:40–47.

56. Li X, et al. Critical role of guanylate binding protein 5 in tumor immune microenvironment and predictive value of immunotherapy response. Front Genet. 2022;13. 10.3389/fgene.2022.984615.

57. Xu D-M, et al. Decoding the impact of MMP1+ malignant subsets on tumor-immune interactions: insights from single-cell and spatial transcriptomics. Cell Death Discov. 2025;11(1):244.

58. Foley CJ, et al. Matrix Metalloprotease-1a Promotes Tumorigenesis and Metastasis*. Journal of Biological Chemistry. 2012;287(29):24330–24338.

59. Pidugu VK, et al. Emerging Functions of Human IFIT Proteins in Cancer. Front Mol Biosci. 2019;6:148.

60. Brombin A, et al. Tfap2b specifies an embryonic melanocyte stem cell that retains adult multifate potential. Cell Rep. 2022;38(2):110234.

61. Brombin A, Patton EE. Melanocyte lineage dynamics in development, growth and disease. Development. 2024;151(15):dev201266.

62. Bueschbell B, Manga P, Schiedel AC. The Many Faces of G Protein-Coupled Receptor 143, an Atypical Intracellular Receptor. Front Mol Biosci. 2022;9. 10.3389/fmolb.2022.873777.

63. Colombo S, et al. Transcriptomic Analysis of Mouse Embryonic Skin Cells Reveals Previously Unreported Genes Expressed in Melanoblasts. Journal of Investigative Dermatology. 2012;132(1):170–178.

64. Chao CC-K, Chang P-Y, Lu HH-P. Human Gas7 isoforms homologous to mouse transcripts differentially induce neurite outgrowth. J Neurosci Res. 2005;81(2):153–162.

65. Lu D, et al. The Tumor-Suppressive Function of UNC5D and Its Repressed Expression in Renal Cell Carcinoma. Clin Cancer Res. 2013;19(11):2883–2892.

66. Kessler D, et al. Targeting Son of Sevenless 1: The pacemaker of KRAS. Current Opinion in Chemical Biology. 2021;62:109–118.

67. Holla VR, et al. ALK: a tyrosine kinase target for cancer therapy. Cold Spring Harb Mol Case Stud. 2017;3(1):a001115.

68. Chang M-M, et al. FGF9/FGFR1 promotes cell proliferation, epithelial-mesenchymal transition, M2 macrophage infiltration and liver metastasis of lung cancer. Translational Oncology. 2021;14(11):101208.

69. Yang D, et al. RasGRP3, a Ras activator, contributes to signaling and the tumorigenic phenotype in human melanoma. Oncogene. 2011;30(45):4590–4600.

70. Hu M, et al. The classification of melanocytic gene signatures. Pigment Cell & Melanoma Research. 2024;37(6):854–863.

71. Li H, Hou L. Regulation of melanocyte stem cell behavior by the niche microenvironment. Pigment Cell & Melanoma Research. 2018;31(5):556–569.

72. De Raeve LE, et al. Distinct phenotypic changes between the superficial and deep component of giant congenital melanocytic naevi: a rationale for curettage. Br J Dermatol. 2006;154(3):485–492.

73. Fujito H, et al. A Case of a Giant Congenital Melanocytic Nevus Treated by Curettage with the Application of Cultured Epidermal Autografts before 6 Months of Age. Plast Reconstr Surg Glob Open. 2021;9(5):e3600.

74. Carney BC, et al. Hypopigmented burn hypertrophic scar contains melanocytes that can be signaled to re-pigment by synthetic alpha-melanocyte stimulating hormone in vitro. PLoS One. 2021;16(3):e0248985.

75. Berndt JD, Halloran MC. Semaphorin 3d promotes cell proliferation and neural crest cell development downstream of TCF in the zebrafish hindbrain. Development. 2006;133(20):3983–3992.

76. Kodo K, et al. Regulation of Sema3c and the Interaction between Cardiac Neural Crest and Second Heart Field during Outflow Tract Development. Sci Rep. 2017;7(1):6771.

77. Gaur P, et al. Role of Class 3 Semaphorins and Their Receptors in Tumor Growth and Angiogenesis. Clin Cancer Res. 2009;15(22):6763–6770.

78. Hao J, et al. Sema3C signaling is an alternative activator of the canonical WNT pathway in glioblastoma. Nat Commun. 2023;14(1):2262.

79. Tam KJ, et al. Semaphorin 3 C drives epithelial-to-mesenchymal transition, invasiveness, and stem-like characteristics in prostate cells. Sci Rep. 2017;7(1):11501.

80. Li S, et al. SEMA3C promotes thyroid cancer via the Wnt/β-catenin pathway. Experimental Cell Research. 2025;444(2):114378.

81. Hu C, et al. Dissecting Melanoma Ecosystem Heterogeneity from Molecular Characteristics to Genetic Variation at Single-Cell Resolution. International Journal of Molecular Sciences. 2025;26(20). 10.3390/ijms26209956.

82. Salehitabar E, et al. Identification of genes with high heterogeneity of expression as a predictor of different prognosis and therapeutic responses in colorectal cancer: a challenge and a strategy. Cancer Cell Int. 2022;22(1):276.

83. Mroz EA, et al. High intratumor genetic heterogeneity is related to worse outcome in patients with head and neck squamous cell carcinoma. Cancer. 2013;119(16):3034–3042.

84. MacDonald WJ, et al. Heterogeneity in Cancer. Cancers. 2025;17(3). 10.3390/cancers17030441.

85. Solé -Boldo L, et al. Single-cell transcriptomes of the human skin reveal age-related loss of fibroblast priming. Commun Biol. 2020;3(1):188.

86. Hao Y, et al. Dictionary learning for integrative, multimodal and scalable single-cell analysis. Nat Biotechnol. 2024;42(2):293–304.

87. Choudhary S, Satija R. Comparison and evaluation of statistical error models for scRNA-seq. Genome Biol. 2022;23(1):27.

88. Yu G, et al. clusterProfiler: an R Package for Comparing Biological Themes Among Gene Clusters. OMICS. 2012;16(5):284–287.

89. Xu S, et al. Using clusterProfiler to characterize multiomics data. Nat Protoc. 2024;19(11):3292–3320.

90. Zhou Z, et al. Spatial transcriptomics deconvolution at single-cell resolution using Redeconve. Nat Commun. 2023;14(1):7930.

